## Supplementary material for "Relating GPCR domains with functionality: receptor helix-bundle and C-terminus differentially influence GRK-specific functions and β-arrestin-mediated regulation": Matthees et al b2V2

Supplementary material
Including Suppl. Fig. 1-11 and Suppl. Tab. 1-6, as well as the respective legends

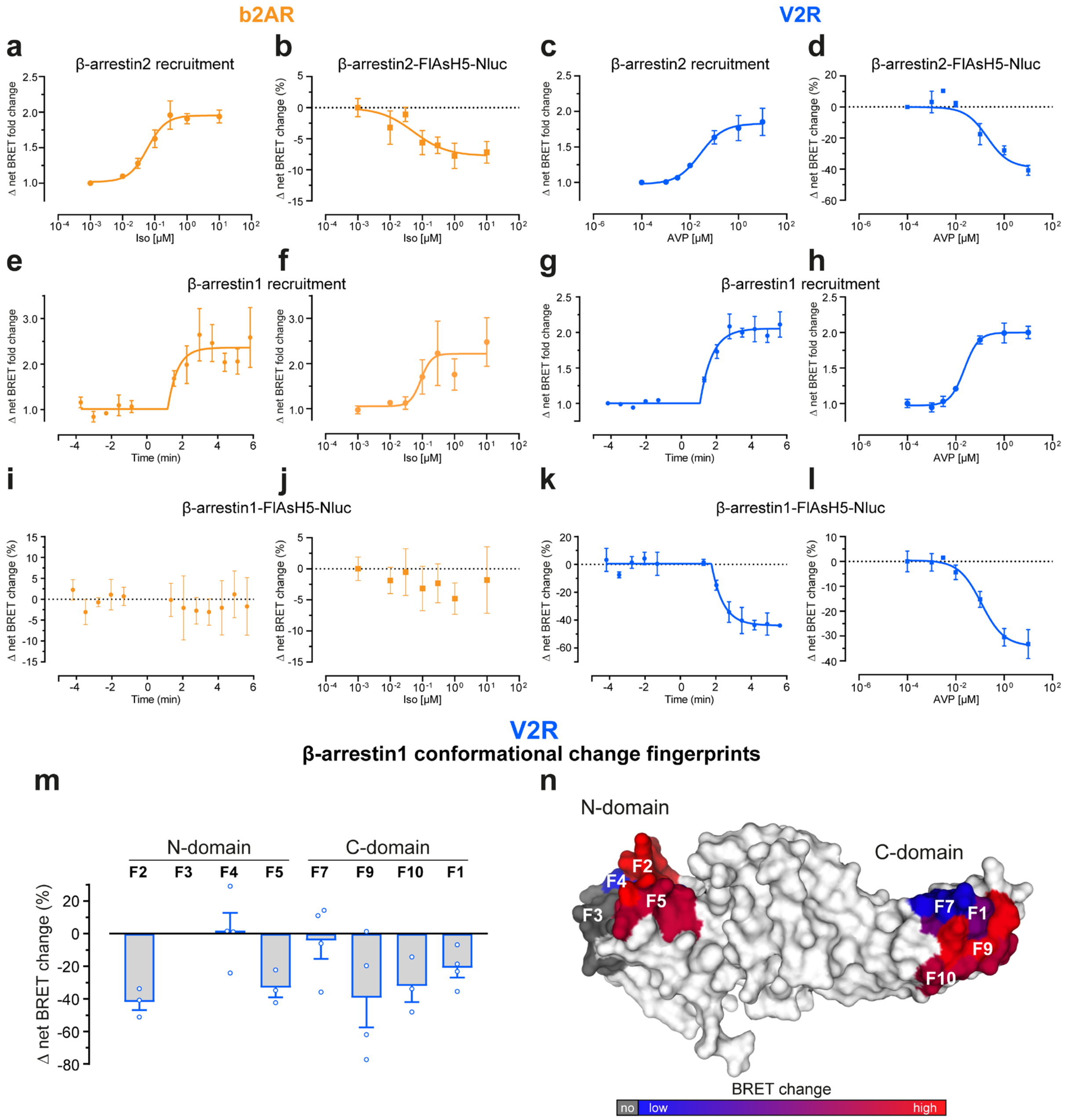


**Supplementary Figure 1: b2AR and V2R differentially interact with β-arrestin1 and 2. a-d,** β-arrestin2 recruitment to b2AR (**a**) and V2R (**c**, data initially published in Drube *et al.*[26]), as well as conformational change of β-arrestin2-FlAsH (F) 5-Nluc in Control cells, co-transfected with untagged b2AR (**b**) or V2R (**d**), as concentration-response curves. Concentration-dependent values correspond to data shown in **Fig.1 b, c, e, f**, averaging timepoints between 2-4 min after stimulation with the indicated agonist concentration. **e-l,** β-arrestin1 recruitment to b2AR (**e, f**) and V2R (**g, h**, data initially published in Drube *et al.*[26]), and β-arrestin1-F5-Nluc conformational changes (**i-l**) were measured in Control cells. BRET changes are shown over time (**e, g, i, k**) following stimulation with 10 µM Iso (e, i) or 3 µM AVP (g, k) or in a concentration-dependent manner (**f, h, j, l**). Data were analyzed analogously to measurements with β-arrestin2 (**Fig.1a-f** and **a-d**) and are shown as Δ net BRET fold change (**a, c, e-h**) or Δ net BRET change (%) (**b, d, i-l**) ± SEM of at least *n=3* independent experiments. **m, n,** Fingerprint of β-arrestin1 conformational change sensors measured in Control cells in presence of untagged V2R. The Δ net BRET changes at 3 µM AVP are shown as bar graphs (**m**). Sensor conditions, which did not fulfill the set pharmacological parameters (Hill slope and $EC_{50}$ analysis as described in methods and **Suppl. Fig. 2**) were classified as non-responding positions and assigned zero. These data were projected onto the surface of the inactive β-arrestin1 crystal structure (PDB: 2WTR) by coloring the respective loop (-fragments) of the labeled FlAsH positions (**n**) ranging from blue to red. The Δ net BRET change was normalized to the maximum sensor signal (red). Non-responding conditions are shown in gray.

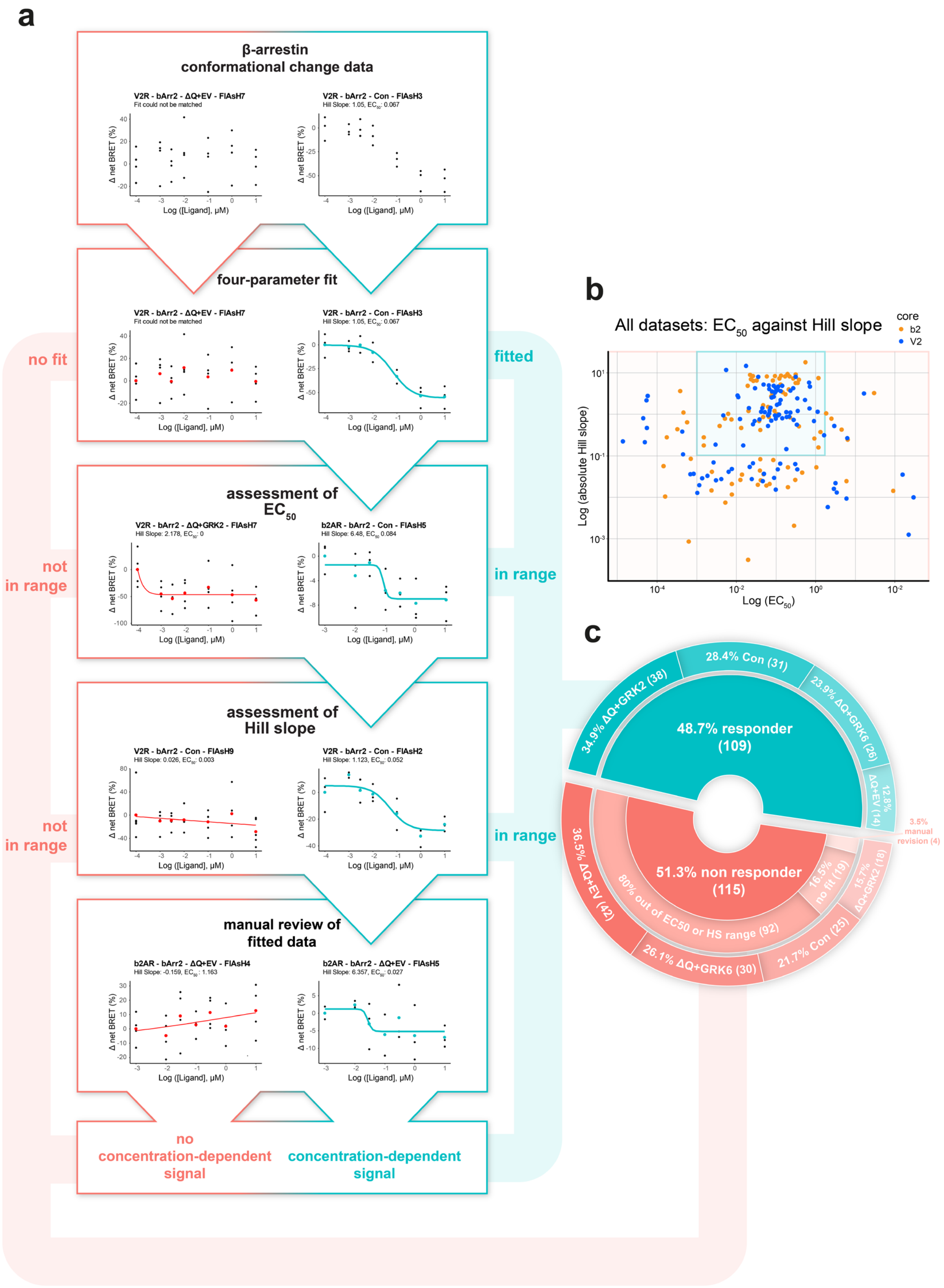


**Supplementary Figure 2: Pharmacological analysis route for the interpretation of biosensor signals. a,** Flowchart detailing individual steps of data analysis to identify “responding” and “non-responding” β-arrestin conformational change signals (featuring example datasets). In brief, concentration-dependent data of at least *n=3* independent experiments served as the input and were fitted using a non-linear four parameter model. Subsequently, fitting parameters ($EC_{50}$ and Hill slope) were assessed and characterized according to set constraints

(Hill slope absolute value > 0.1, $EC_{50}$ between $10^{-3}$ and $10^{0.3}$ μM). Finally, all datasets were subjected to manual review. Data for which a four-parameter fit was not applicable and/or did not adhere to set constraints were classified as “non-responding” and hence interpreted as 0% conformational change throughout the study. **b,** Scatterplot showing $EC_{50}$ and absolute Hill slope values of all measured conditions. Set constraints used to identify “responding” β-arrestin conformational change signals are indicated with a green box. **c,** Donut plot that shows the makeup and characterization of all β-arrestin conformational change data included in this study, detailing the exact reasons for “non-responder” identification, as well as the individual fractions of cellular GRK-specific conditions.

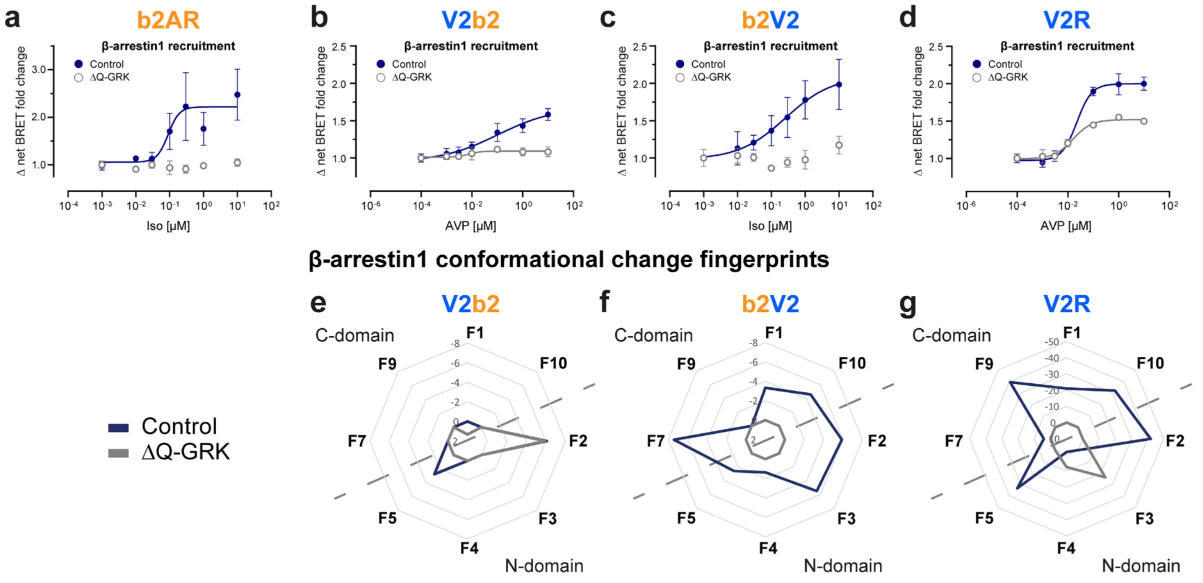


**Supplementary Figure 3: Different combinations of GPCR helix bundles and C-termini orchestrate distinct, GRK-dependent β-arrestin1 interactions and conformational changes. a-d,** β-arrestin1 recruitment to the b2AR (**a**), V2b2 (**b**), b2V2 (**c**) and V2R (**d**) in ΔQ-GRK or Control cells stimulated with Iso (**a, c**) or AVP (**b, d**) as indicated. Data for b2AR, b2V2 and V2R were initially published in Drube *et al.*[26] and are shown again to allow a direct comparison. Data are shown as Δ net BRET fold change over baseline ± SEM of *n*=3 independent experiments. **e-g,** Complete β-arrestin1 conformational change fingerprints are shown for each receptor in Control and ΔQ-GRK cells, stimulated with 10 μM Iso (**f**) or 3 μM AVP (**e, g**), as Δ net BRET change (%) as radar plots, analogously to the data for β-arrestin2 (**Fig. 2s-v**). Sensor conditions, which did not fulfill the pharmacological parameters (Hill slope and $EC_{50}$ analysis as described in methods and **Suppl. Fig. 2**) were classified as non-responding conditions and assigned zero.

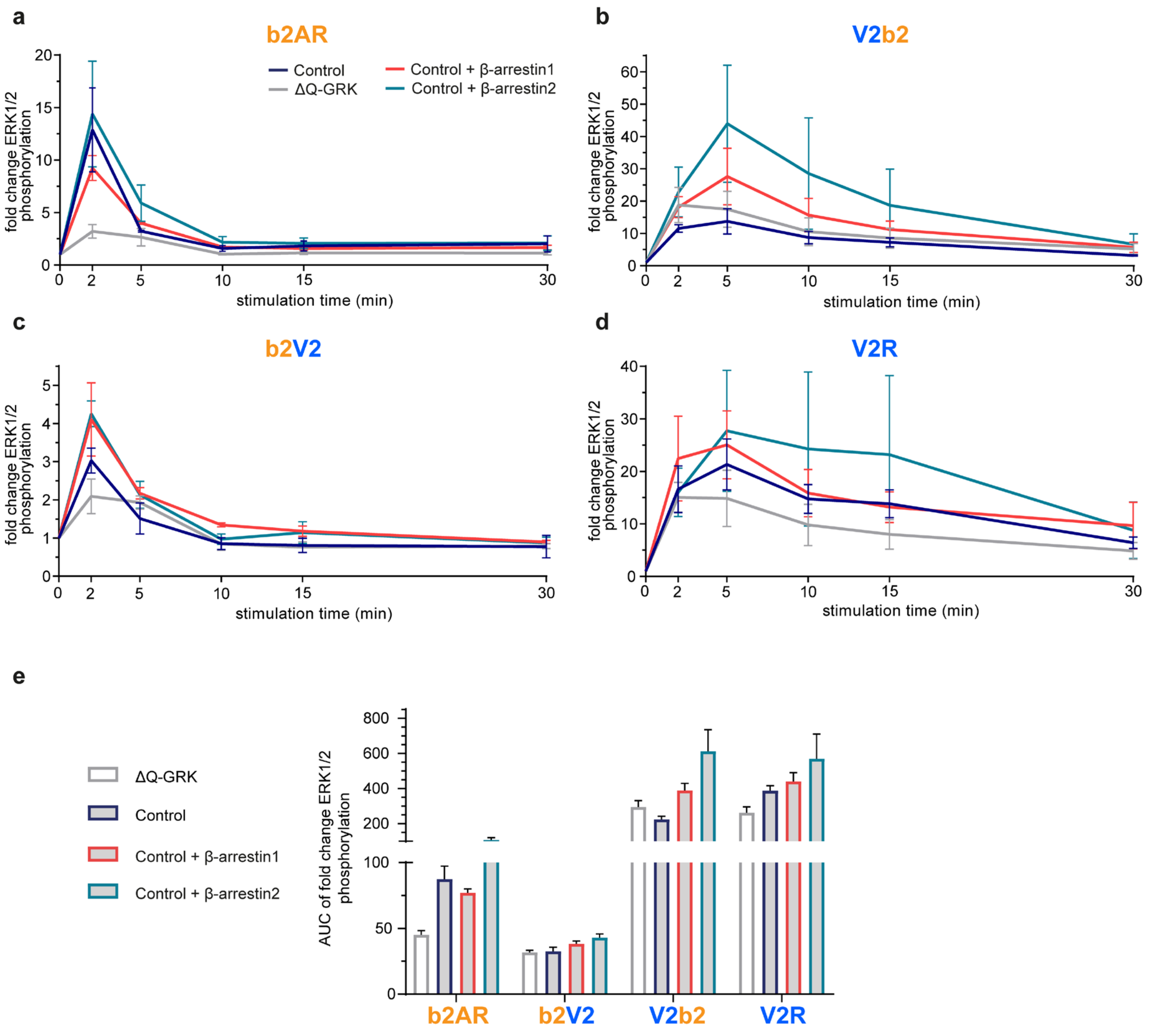


**Supplementary Figure 4: Overexpression of β-arrestin enhances ERK1/2 phosphorylation. a-d,** Western blots detecting phosphorylated ERK (pERK) and total ERK over time in Control or ΔQ-GRK cells, stably expressing b2AR (**a**), V2b2 (**b**), b2V2 (**c**) or V2R (**d**), of *n=3* independent experiments were quantified. Data are normalized to the respective basal levels of pERK and are shown as fold change ERK1/2 phosphorylation over time ± SEM. Measurements in Control and ΔQ-GRK cells under endogenous expression of β-arrestins from **Fig. 3e-h** are shown again for each receptor construct to enable direct comparison. **e,** As displayed in **Fig. 3e-h**, the area under the curve (AUC) was quantified for each condition to compare the pERK changes over time, shown as a bar graph. Statistical significance was analyzed using a 2-way ANOVA. Complete results of the statistical analysis can be accessed in **Suppl. Tab. 1**.

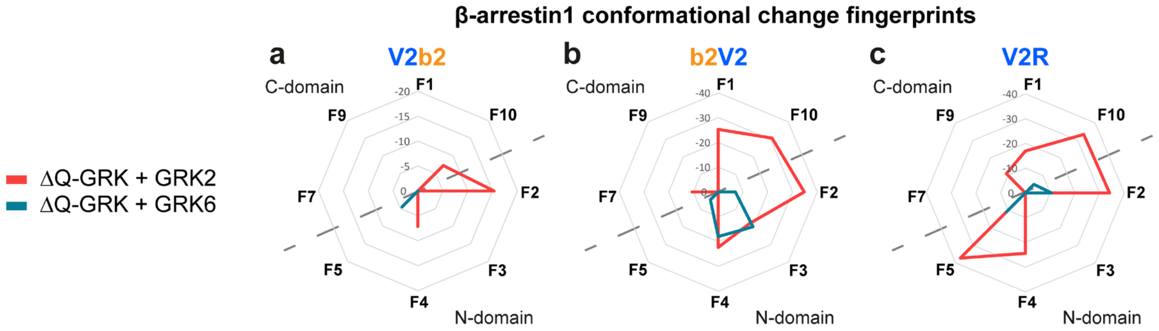


**Supplementary Figure 5: Differential GPCR phosphorylation by GRK2 and GRK6 induces distinct receptor-specific β-arrestin1 conformational changes. a-c,** Fingerprint of β-arrestin1 conformational change sensors measured in ΔQ-GRK cells, individually overexpressing GRK2 or GRK6, in presence of untagged V2b2 (**a**), b2V2 (**b**) or V2R (**c**). The Δ net BRET changes at 10 µM Iso (**b**) or 3 µM AVP (**a, c**) are shown as Δ net BRET change (%) of at least *n=3* independent experiments as radar plots, analogously to **Fig. 4i-l**. Sensor conditions, which did not fulfill the set pharmacological parameters (Hill slope and $EC_{50}$ analysis as described in methods) were classified as non-responding conditions and assigned zero.

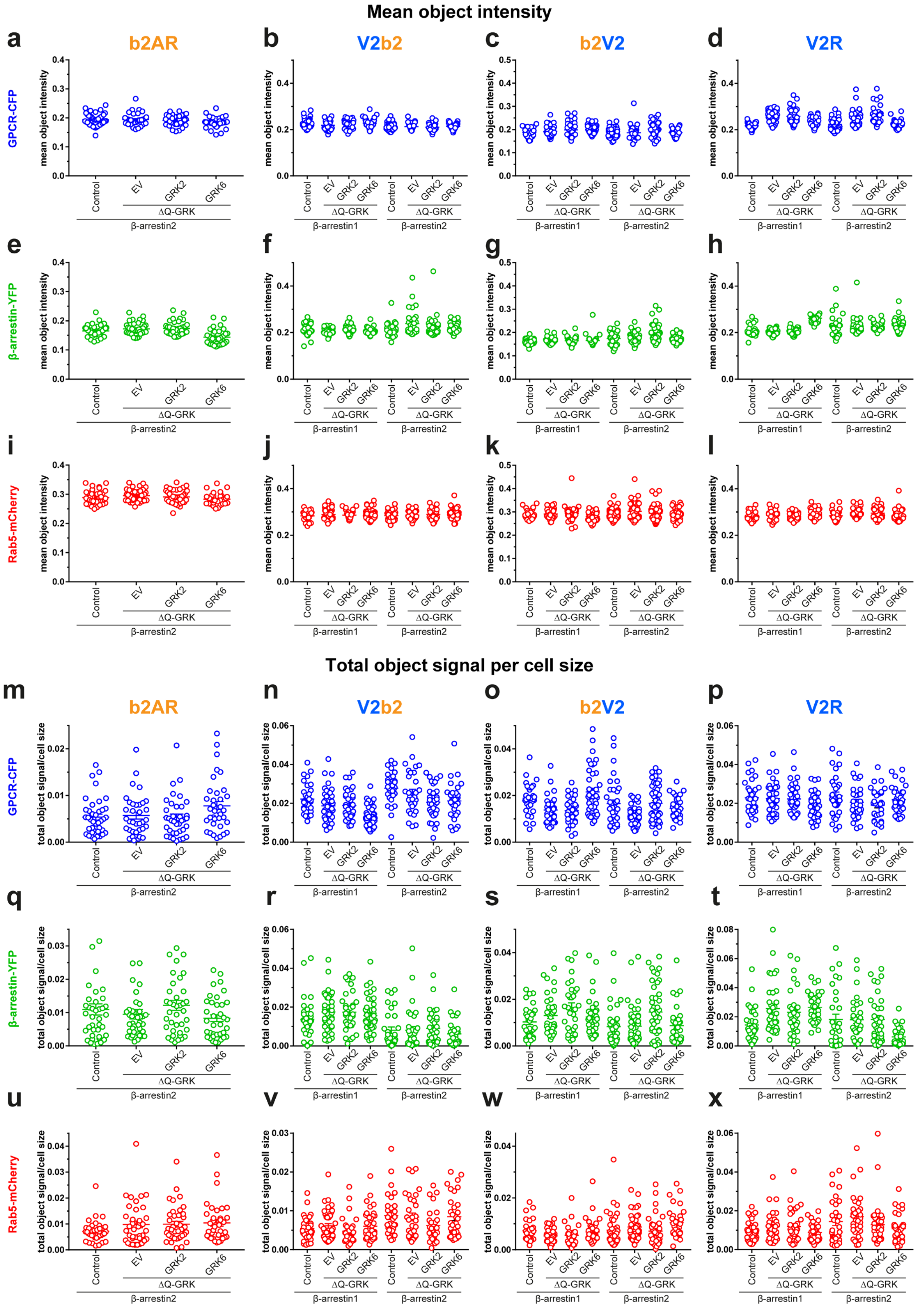


**Supplementary Figure 6: Overview of mean object intensity and total object signal per cell size for each microscopy condition.** Squassh segmentation parameters (mean object intensity (**a-l**) and total object per cell size (**m-x**)) for all non-stimulated confocal images that were subjected to co-localization analysis, separated by channels (GPCR-CFP (blue), β-arrestin-YFP (green), Rab5-mCherry (red)).

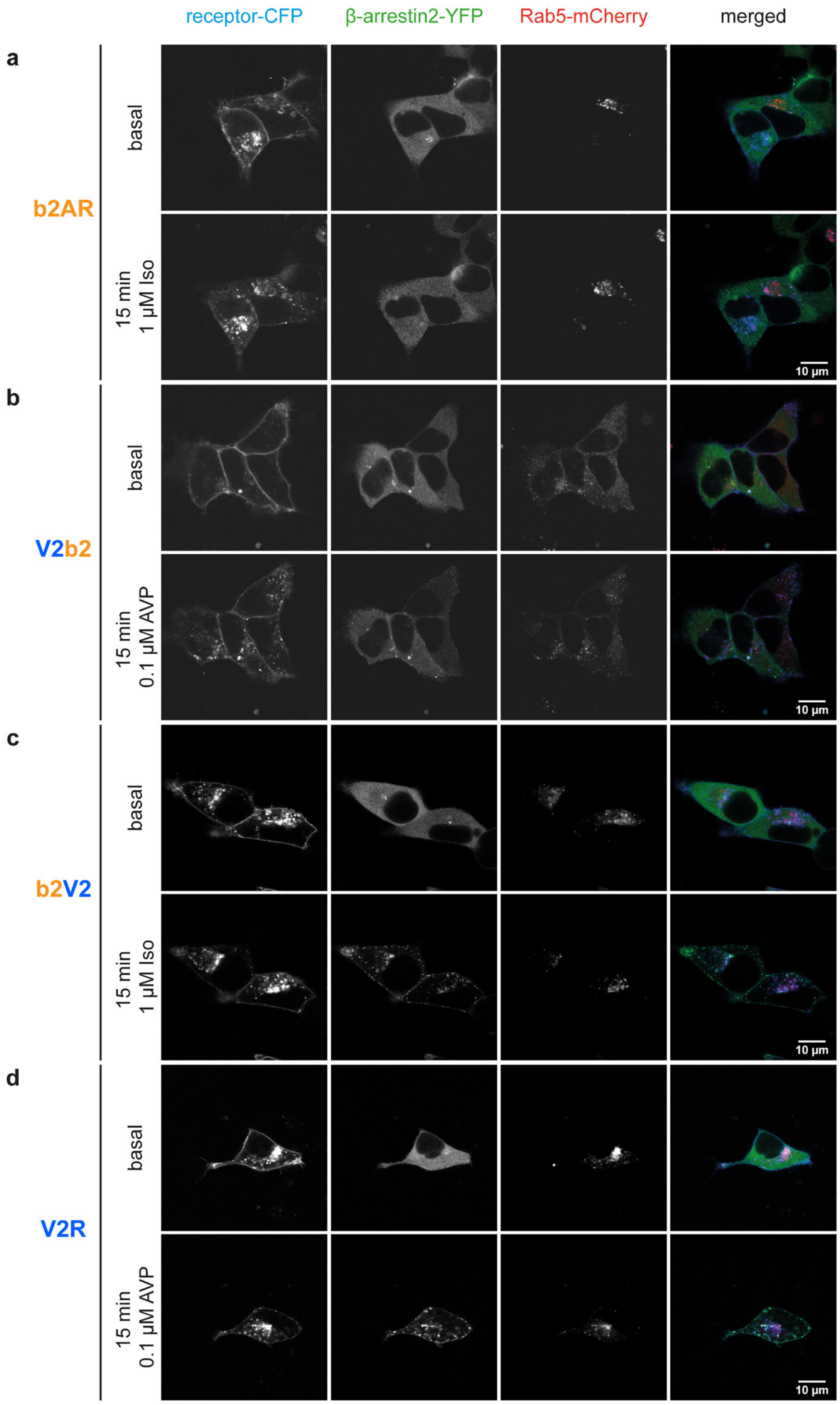


**Supplementary Figure 7: Single channel images show β-arrestin translocation to early endosomes depends on the receptor C-terminus. a-d,** Control cells were transfected with the indicated receptor-CFP (blue), β-arrestin-YFP (green) and early endosome marker Rab5-mCherry (red). Confocal images were taken before (basal) and after 15 min stimulation with the specified ligand (stimulated). Here, single channel and merged images are displayed for all receptors of the representative Control cell images shown in **Fig. 5a**.

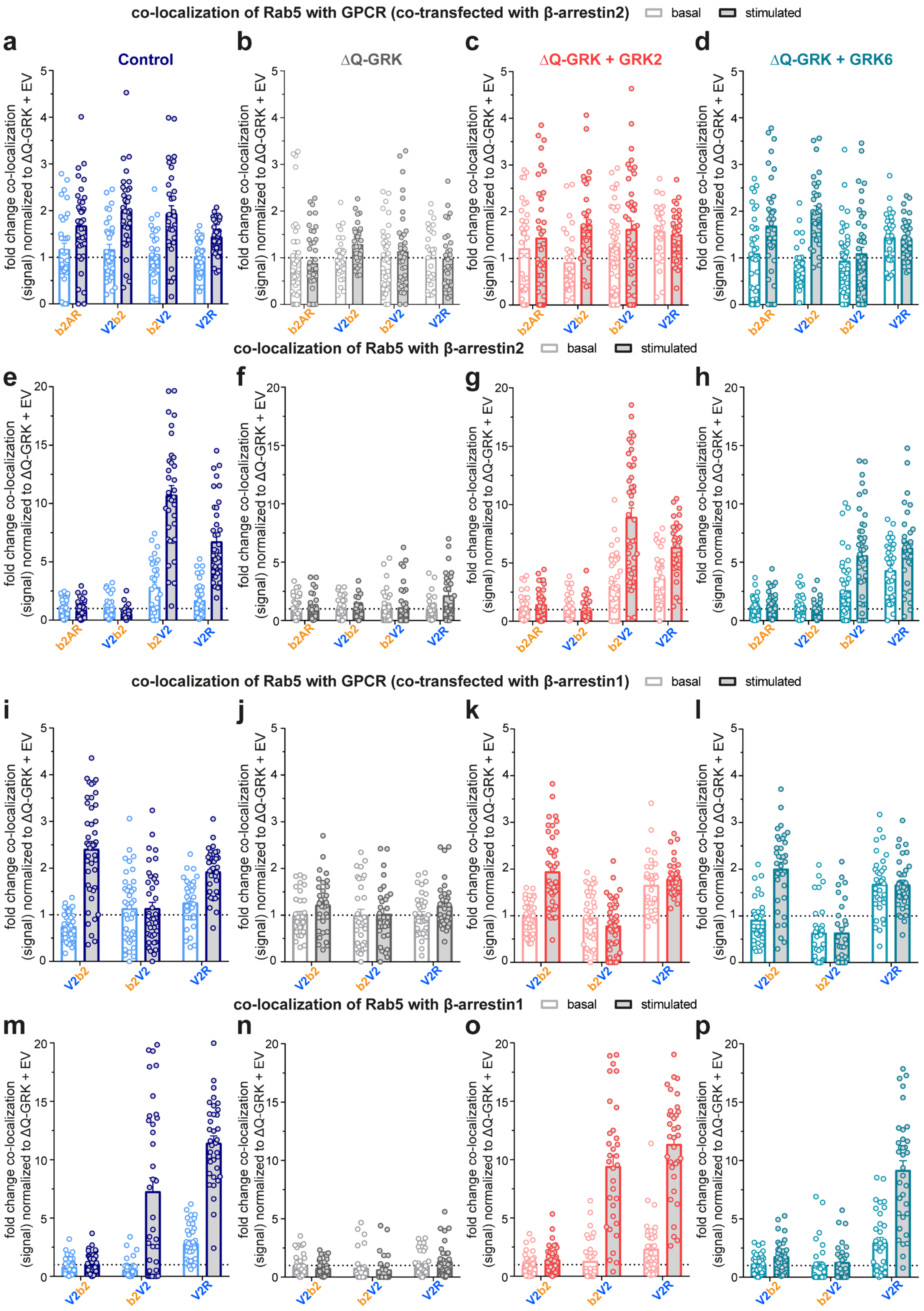


**Supplementary Figure 8: Quantification of receptor and β-arrestin translocation to early endosomes. a-h,** Control or ΔQ-GRK cells were transfected with the indicated receptor-CFP, β-arrestin2-YFP and early endosome marker Rab5-mCherry, as well as GRK2 or GRK6 in ΔQ-GRK cells, as indicated. Confocal images were taken before (basal; lighter color) and after 15 min of stimulation with the specified ligand (stimulated; darker color). The co-localization (signal) of the receptor with Rab5 in presence of β-arrestin2 (**a-d**) or of β-arrestin2 with Rab 5 for each receptor construct (**e-h**) was quantified using Squassh and SquasshAnalyst[44,45]. The co-localization of Rab5 with β-arrestin2 in Control cells (**Fig. 5b**) is displayed again (**e**) to allow a direct comparison between the measured GRK conditions. **i-p,** Analogously to the experiments performed with β-arrestin2, co-localization with Rab5 was also investigated in presence of β-arrestin1-YFP for V2b2, b2V2 and V2R. For all conditions, data of at least 28 images per condition are shown as mean fold change in co-localization (signal) + SEM, normalized to the respective unstimulated (basal) ΔQ-GRK + EV condition. Statistical comparison between basal and stimulated values was performed using a two-way ANOVA, followed by a Sidak's test. Basal and stimulated values between different conditions were compared with a two-way ANOVA, followed by a Tukey's test. Detailed results can be accessed in **Suppl. Tab. 3-6**.

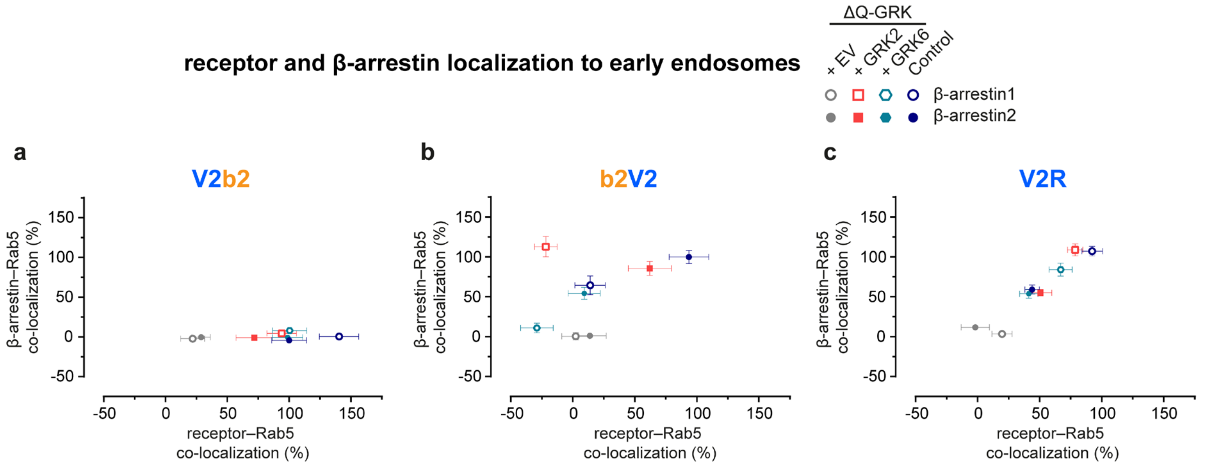


**Supplementary Figure 9: β-arrestin1 translocation to early endosomes is controlled by the receptor C-terminus. a-c,** Analogous to the data shown in **Fig. 5d-f**, the fold change in co-localization (signal) was analyzed for the receptor or β-arrestin1 with Rab5, as fold change over the respective basal ΔQ-GRK + EV condition for each tested receptor (V2b2 (**a**), b2V2 (**b**), V2R (**c**)) and GRK condition (Control, ΔQ-GRK +EV, +GRK2, +GRK6). The stimulated values for receptor–Rab5 co-localization are show on the x-axis and for β-arrestin1–Rab5 co-localization on the y-axis. For both dimension, the data points were normalized to the respective maximum in presence of β-arrestin2 (for receptor–Rab5 co-localization: V2b2 in Control cells, for β-arrestin2–Rab5 co-localization: b2V2 in Control cells) and are shown in percent. Data for β-arrestin2 (**Fig. 5c-f**) are shown again to allow a direct comparison. Detailed statistical analysis can be accessed in **Suppl. Tab. 4-6**.

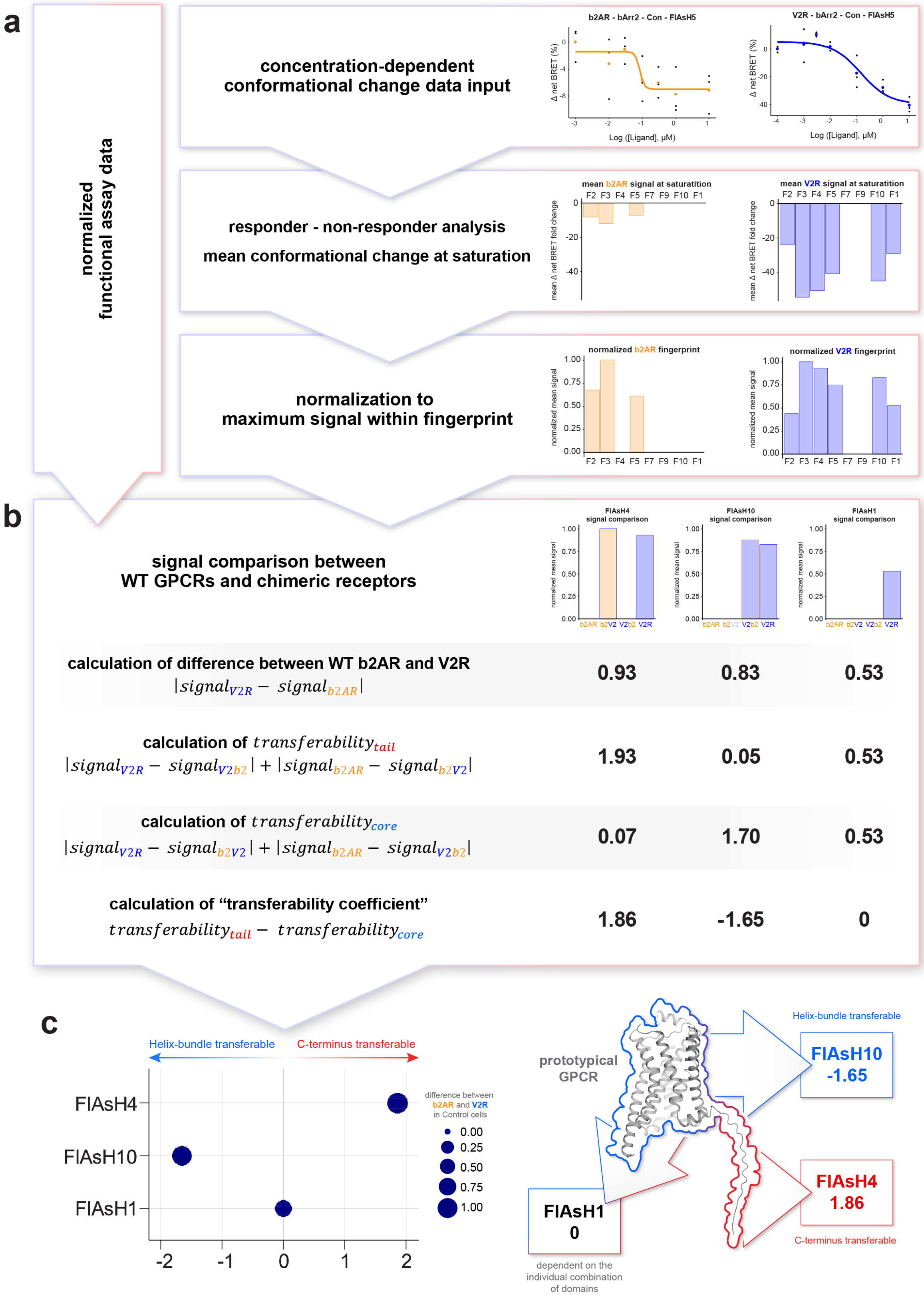


**Supplementary Figure 10: Influence of helix-bundle or C-terminus on β-arrestin conformational change signals, calculated via “transferability coefficients”. a,** Flowchart indicating normalization steps of input β-arrestin conformational change and functional assay data (featuring example data on the right). **b,** Calculation of “transferability coefficients” using normalized β-arrestin conformational change signals and normalized functional assay data at saturating ligand concentrations for all four receptor specific conditions (b2AR, b2V2, V2b2 and V2R, featuring example data for β-arrestin conformational changes on the right). First, the difference between WT GPCR-induced signals was calculated, followed by the assessment of differences between receptor-specific conditions that share the same helix-bundle ($\text{transferability}_{\text{tail}}$) and C-terminus ($\text{transferability}_{\text{core}}$). Finally, subtraction of calculated $\text{transferability}_{\text{tail}}$ and $\text{transferability}_{\text{core}}$ values yields unitless transferability coefficients. A negative coefficient indicates a more pronounced influence of the helix-bundle, whereas a positive coefficient is interpreted as the GPCR C-terminus having a more pronounced influence on the molecular movement of individual β-arrestin conformational change biosensors. A value of zero suggests that the receptor-specific signals depend on the individual combination of domains. **c,** Calculated transferability coefficients of β-arrestin2-F1, F4 and F10 biosensors in Control cells (used as example data above) are plotted to indicate the influence of individual GPCR domains on conformational changes of these specific β-arrestin2 moieties. The difference in signal between WT GPCRs is shown as the size of the plotted symbols.

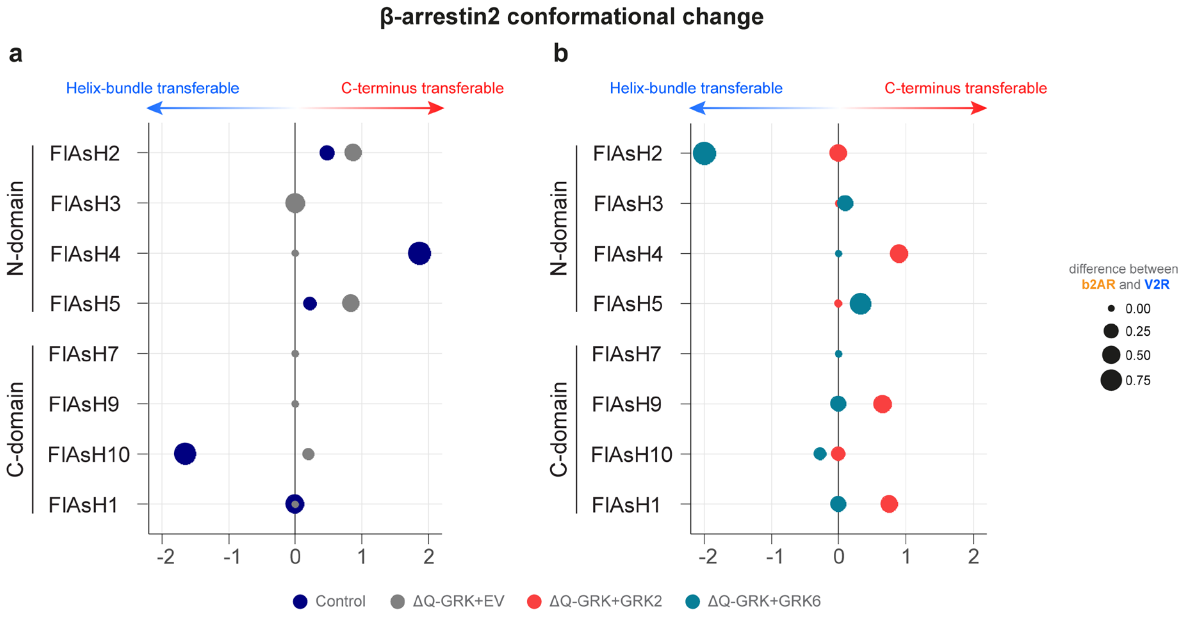


**Supplementary Figure 11: "Transferability coefficents" of GRK-specific β-arrestin2 conformational changes. a, b,** Calculated transferability coefficients of β-arrestin2 biosensors in Control cells or EV-transfected ΔQ-GRK cells (**a**) and under GRK2 or 6 overexpression (**b**) are plotted to indicate the influence of individual GPCR domains on conformational changes. The difference in signal between WT GPCRs is shown as the size of the plotted symbols. Calculation of these coefficients is explained in **Suppl. Fig. 10**.

**Supplementary Table 1:** Detailed statistical results of the analysis of the area under the curve (AUC), derived from Western blots, shown in **Fig. 3e-h** and **Suppl. Fig. 4e**. AUC of fold change of ERK1/2 phosphorylation over time was compared between Control and ΔQ-GRK cells, stably expressing the b2AR, V2b2, b2V2 or V2R, using an unpaired, two-tailed t-test (**Fig. 3e-h**) or between Control cells in absence or presence of transient overexpression of β-arrestin1 or β-arrestin2 with a two-way ANOVA, followed by a Dunnett's test (**Suppl. Fig. 4e**). For each condition the mean difference, 95% confidence interval (CI) of the difference, and the adjusted *p* value are shown (*ns,* not significant; *, $p < 0.05$; **, $p < 0.01$; ***, $p < 0.001$; ****, $p < 0.0001$).

| **conditions** | | | | |
|---|---|---|---|---|
| **AUC of pERK over time** | **unpaired two-tailed t-test** | | | |
| Control vs ΔQ-GRK | **diff. betw. means** | **95% CI** | ***p* value** | |
| b2AR | 42.34 ± 10.52 | 13.14 to 71.54 | 0.0158 | * |
| V2b2 | -71.10 ± 40.74 | -184.2 to 42.00 | 0.1559 | ns |
| b2V2 | 1.050 ± 3.426 | -8.461 to 10.56 | 0.7745 | ns |
| V2R | 124.4 ± 44.47 | 0.9307 to 247.9 | 0.0489 | * |
| **AUC of pERK over time** | **two-way ANOVA, followed by a Dunnetts's test** | | | |
| Control cells + EV vs. + β-arrestin1 | **mean diff.** | **95% CI of diff.** | **adjusted *p* value** | |
| b2AR | 10.37 | -17.84 to 38.58 | 0.5713 | ns |
| V2b2 | -165.3 | -390.4 to 59.80 | 0.1569 | ns |
| b2V2 | -5.54 | -14.74 to 3.657 | 0.2632 | ns |
| V2R | -53.4 | -291.7 to 184.9 | 0.8015 | ns |
| Control cells + EV vs. + β-arrestin2 | | | | |
| b2AR | -20.65 | -48.86 to 7.565 | 0.1585 | ns |
| V2b2 | -388.4 | -613.5 to -163.3 | 0.0019 | ** |
| b2V2 | -10.21 | -19.41 to -1.013 | 0.0304 | * |
| V2R | -183.3 | -421.6 to 55.01 | 0.1359 | ns |

**Supplementary Table 2:** Detailed statistical results of the analysis of the normalized OD 405 nm (%) for the proximal and distal phosphorylation cluster in ΔQ-GRK cells stably overexpressing one of the investigated receptors and GRK2 or 6 in comparison to the respective Control cells (**Fig. 4a-h**). Values of maximal ligand concentration and vehicle stimulation were compared using a two-way ANOVA, followed by a Tukey's test. For each condition the mean difference, 95% confidence interval (CI) of the difference, and the adjusted *p* value are shown (*ns,* not significant; *, $p < 0.05$; **, $p < 0.01$; ***, $p < 0.001$; ****, $p < 0.0001$).

| conditions | two-way ANOVA, followed by Tukey's test | | | |
|---|---|---|---|---|
| normalized OD 405 nm | | | | |
| b2AR - proximal phosphorylation | mean diff. | 95% CI of diff. | adjusted *p* value | |
| vehicle stimulation | | | | |
| Con vs. GRK2 | -4.938 | -13.99 to 4.115 | 0.3761 | ns |
| Con vs. GRK6 | -16.3 | -25.35 to -7.245 | 0.0004 | *** |
| GRK2 vs. GRK6 | -11.36 | -20.41 to -2.307 | 0.0121 | * |
| maximal stimulation | | | | |
| Con vs. GRK2 | 59.7 | 50.65 to 68.76 | <0.0001 | **** |
| Con vs. GRK6 | -7.346 | -16.4 to 1.707 | 0.1275 | ns |
| GRK2 vs. GRK6 | -67.05 | -76.1 to -58 | <0.0001 | **** |
| b2AR - distal phosphorylation | | | | |
| vehicle stimulation | | | | |
| Con vs. GRK2 | -1.796 | -19.61 to 16.02 | 0.9657 | ns |
| Con vs. GRK6 | -5.46 | -23.28 to 12.36 | 0.7275 | ns |
| GRK2 vs. GRK6 | -3.664 | -21.48 to 14.15 | 0.8655 | ns |
| maximal stimulation | | | | |
| Con vs. GRK2 | 30.1 | 12.28 to 47.92 | 0.0009 | *** |
| Con vs. GRK6 | -2.344 | -20.16 to 15.47 | 0.9424 | ns |
| GRK2 vs. GRK6 | -32.44 | -50.26 to -14.62 | 0.0004 | *** |
| V2b2 - proximal phosphorylation | | | | |
| vehicle stimulation | | | | |
| Con vs. GRK2 | -0.408 | -13.11 to 12.29 | 0.9965 | ns |
| Con vs. GRK6 | -11.48 | -24.18 to 1.22 | 0.0818 | ns |
| GRK2 vs. GRK6 | -11.07 | -23.78 to 1.628 | 0.0957 | ns |
| maximal stimulation | | | | |
| Con vs. GRK2 | 64.07 | 51.36 to 76.77 | <0.0001 | **** |
| Con vs. GRK6 | 32 | 19.3 to 44.7 | <0.0001 | **** |
| GRK2 vs. GRK6 | -32.07 | -44.77 to -19.37 | <0.0001 | **** |
| V2b2 - distal phosphorylation | | | | |
| vehicle stimulation | | | | |
| Con vs. GRK2 | 0.122 | -8.521 to 8.765 | 0.9993 | ns |
| Con vs. GRK6 | -5.258 | -13.9 to 3.385 | 0.2999 | ns |
| GRK2 vs. GRK6 | -5.38 | -14.02 to 3.263 | 0.2843 | ns |
| maximal stimulation | | | | |
| Con vs. GRK2 | 55.16 | 46.52 to 63.8 | <0.0001 | **** |
| Con vs. GRK6 | 42.98 | 34.34 to 51.62 | <0.0001 | **** |
| GRK2 vs. GRK6 | -12.18 | -20.83 to -3.541 | 0.0048 | ** |
| b2V2 - proximal phosphorylation | | | | |
| vehicle stimulation | | | | |
| Con vs. GRK2 | -1.938 | -21.66 to 17.78 | 0.9674 | ns |
| Con vs. GRK6 | -5.334 | -25.06 to 14.39 | 0.7799 | ns |
| GRK2 vs. GRK6 | -3.396 | -23.12 to 16.33 | 0.9035 | ns |
| maximal stimulation | | | | |
| Con vs. GRK2 | 59.28 | 39.56 to 79.01 | <0.0001 | **** |
| Con vs. GRK6 | 12.46 | -7.267 to 32.18 | 0.2746 | ns |
| GRK2 vs. GRK6 | -46.83 | -66.55 to -27.11 | <0.0001 | **** |
| b2V2 - distal phosphorylation | | | | |
| vehicle stimulation | | | | |
| Con vs. GRK2 | -4.404 | -29.52 to 20.71 | 0.9001 | ns |
| Con vs. GRK6 | -16.05 | -41.16 to 9.061 | 0.2666 | ns |
| GRK2 vs. GRK6 | -11.65 | -36.76 to 13.46 | 0.4888 | ns |
| maximal stimulation | | | | |
| Con vs. GRK2 | 59.66 | 34.54 to 84.77 | <0.0001 | **** |
| Con vs. GRK6 | 1.434 | -23.68 to 26.55 | 0.9889 | ns |
| GRK2 vs. GRK6 | -58.22 | -83.33 to -33.11 | <0.0001 | **** |
| V2R - proximal phosphorylation | | | | |
| vehicle stimulation | | | | |
| Con vs. GRK2 | 1.108 | -11.58 to 13.8 | 0.9742 | ns |
| Con vs. GRK6 | -13.25 | -25.94 to -0.5594 | 0.0395 | * |
| GRK2 vs. GRK6 | -14.36 | -27.05 to -1.667 | 0.0245 | * |
| maximal stimulation | | | | |
| Con vs. GRK2 | 43.88 | 31.19 to 56.57 | <0.0001 | **** |
| Con vs. GRK6 | 17.67 | 4.975 to 30.36 | 0.0053 | ** |
| GRK2 vs. GRK6 | -26.22 | -38.91 to -13.53 | <0.0001 | **** |
| V2R - distal phosphorylation | | | | |
| vehicle stimulation | | | | |
| Con vs. GRK2 | 0.058 | -15.67 to 15.79 | >0.9999 | ns |
| Con vs. GRK6 | -12.16 | -27.89 to 3.567 | 0.1518 | ns |
| GRK2 vs. GRK6 | -12.22 | -27.95 to 3.509 | 0.1493 | ns |
| maximal stimulation | | | | |
| Con vs. GRK2 | 42.96 | 27.23 to 58.69 | <0.0001 | **** |
| Con vs. GRK6 | 26.76 | 11.03 to 42.49 | 0.0008 | *** |
| GRK2 vs. GRK6 | -16.2 | -31.93 to -0.4653 | 0.0427 | * |

**Supplementary Table 3:** Detailed statistical results of the analysis of the microscopy co-localization quantification, shown in **Fig. 5b** (for co-localization of Rab5 with β-arrestin2 in Control cells) and **Suppl. Fig. 8**. Here, basal and stimulated values of co-localization for each tested receptor with Rab5, in presence of β-arrestin2 or β-arrestin1, as well as for β-arrestin2 or β-arrestin1 with Rab5 were compared using a two-way ANOVA, followed by a Sidak's test (*ns,* not significant; *, $p < 0.05$; **, $p < 0.01$; ***, $p < 0.001$; ****, $p < 0.0001$). For each condition the mean difference, 95% confidence interval (CI) of the difference, and the adjusted *p* value are shown.

| **conditions** | **two-way ANOVA, followed by Sidak's test** | | | |
|---|---|---|---|---|
| **co-localization of Rab5 with GPCR (β-arrestin2)** | | | | |
| basal vs. stim in Control | **mean diff.** | **95% CI of diff.** | **adjusted *p* value** | |
| b2AR | -0.5116 | -0.923 to -0.1001 | 0.0081 | ** |
| V2b2 | -0.8347 | -1.264 to -0.4051 | <0.0001 | **** |
| b2V2 | -0.8738 | -1.311 to -0.4367 | <0.0001 | **** |
| V2R | -0.4242 | -0.8384 to -0.009991 | 0.0424 | * |
| basal vs. stim in ΔQ-GRK | | | | |
| b2AR | 0.1144 | -0.2837 to 0.5125 | 0.9222 | ns |
| V2b2 | -0.2891 | -0.7269 to 0.1487 | 0.3406 | ns |
| b2V2 | -0.1381 | -0.5362 to 0.2599 | 0.8568 | ns |
| V2R | 0.01865 | -0.4343 to 0.4716 | >0.9999 | ns |
| basal vs. stim in ΔQ-GRK + GRK2 | | | | |
| b2AR | -0.225 | -0.7439 to 0.2939 | 0.7281 | ns |
| V2b2 | -0.8062 | -1.352 to -0.2605 | 0.001 | ** |
| b2V2 | -0.3911 | -0.8749 to 0.09272 | 0.1635 | ns |
| V2R | 0.07107 | -0.4831 to 0.6252 | 0.996 | ns |
| basal vs. stim in ΔQ-GRK cells | | | | |
| b2AR | -0.5955 | -1.029 to -0.1621 | 0.0026 | ** |
| V2b2 | -1.176 | -1.635 to -0.7174 | <0.0001 | **** |
| b2V2 | -0.1591 | -0.5491 to 0.231 | 0.77 | ns |
| V2R | 0.02669 | -0.422 to 0.4754 | 0.9998 | ns |
| **co-localization of Rab5 with GPCR (β-arrestin1)** | | | | |
| basal vs. stim in Control | | | | |
| V2b2 | -1.656 | -2.021 to -1.291 | <0.0001 | **** |
| b2V2 | -0.00115 | -0.3824 to 0.3801 | >0.9999 | ns |
| V2R | -0.6655 | -1.077 to -0.2545 | 0.0004 | *** |
| basal vs. stim in ΔQ-GRK | | | | |
| V2b2 | -0.2202 | -0.5477 to 0.1073 | 0.2879 | ns |
| b2V2 | -0.02428 | -0.3622 to 0.3136 | 0.9974 | ns |
| V2R | -0.1975 | -0.5155 to 0.1206 | 0.3559 | ns |
| basal vs. stim in ΔQ-GRK + GRK2 | | | | |
| V2b2 | -0.9736 | -1.274 to -0.6735 | <0.0001 | **** |
| b2V2 | 0.1774 | -0.1409 to 0.4956 | 0.4512 | ns |
| V2R | -0.1337 | -0.4584 to 0.191 | 0.6897 | ns |
| basal vs. stim in ΔQ-GRK cells | | | | |
| V2b2 | -1.094 | -1.467 to -0.7197 | <0.0001 | **** |
| b2V2 | -0.001348 | -0.4026 to 0.3999 | >0.9999 | ns |
| V2R | 0.009035 | -0.3647 to 0.3828 | >0.9999 | ns |
| **co-localization of Rab5 with β-arrestin2** | | | | |
| basal vs. stim in Control | | | | |
| b2AR | 0.1515 | -1.2 to 1.503 | 0.9976 | ns |
| V2b2 | 0.4558 | -0.9749 to 1.887 | 0.8907 | ns |
| b2V2 | -7.924 | -9.367 to -6.481 | <0.0001 | **** |
| V2R | -5.039 | -6.39 to -3.688 | <0.0001 | **** |
| basal vs. stim in ΔQ-GRK | | | | |
| b2AR | 0.05554 | -0.6855 to 0.7966 | 0.9995 | ns |
| V2b2 | 0.02879 | -0.7982 to 0.8558 | >0.9999 | ns |
| b2V2 | -0.1135 | -0.8645 to 0.6375 | 0.9924 | ns |
| V2R | -1.147 | -1.988 to -0.3063 | 0.0029 | ** |
| basal vs. stim in ΔQ-GRK + GRK2 | | | | |
| b2AR | -0.2852 | -1.753 to 1.182 | 0.9805 | ns |
| V2b2 | 0.245 | -1.352 to 1.842 | 0.992 | ns |
| b2V2 | -5.928 | -7.35 to -4.506 | <0.0001 | **** |
| V2R | -2.64 | -4.277 to -1.004 | 0.0003 | *** |
| basal vs. stim in ΔQ-GRK cells | | | | |
| b2AR | -0.2084 | -1.58 to 1.163 | 0.9923 | ns |
| V2b2 | 0.3268 | -1.159 to 1.812 | 0.9694 | ns |
| b2V2 | -2.999 | -4.296 to -1.702 | <0.0001 | **** |
| V2R | -1.933 | -3.374 to -0.4926 | 0.0035 | ** |
| **co-localization of Rab5 with β-arrestin1** | | | | |
| basal vs. stim in Control | | | | |
| V2b2 | -0.2404 | -1.881 to 1.401 | 0.9792 | ns |
| b2V2 | -6.711 | -8.491 to -4.931 | <0.0001 | **** |
| V2R | -8.609 | -10.44 to -6.779 | <0.0001 | **** |
| basal vs. stim in ΔQ-GRK | | | | |
| V2b2 | 0.222 | -0.4559 to 0.9 | 0.8159 | ns |
| b2V2 | 0.1382 | -0.5789 to 0.8554 | 0.9545 | ns |
| V2R | -0.3441 | -1.002 to 0.3142 | 0.506 | ns |
| basal vs. stim in ΔQ-GRK + GRK2 | | | | |
| V2b2 | -0.4809 | -2.049 to 1.087 | 0.8436 | ns |
| b2V2 | -8.121 | -9.868 to -6.375 | <0.0001 | **** |
| V2R | -8.982 | -10.7 to -7.263 | <0.0001 | **** |
| basal vs. stim in ΔQ-GRK cells | | | | |
| V2b2 | -0.64 | -2.069 to 0.7888 | 0.6299 | ns |
| b2V2 | -0.2647 | -1.773 to 1.243 | 0.965 | ns |
| V2R | -6.207 | -7.656 to -4.757 | <0.0001 | **** |

**Supplementary Table 4:** Detailed statistical results of the analysis of the microscopy co quantification, shown in **Suppl. Fig. 8**. Here, basal or stimulated values of co-localization with Rab were compared between tested receptors, in presence of β-arrestin2, using a two-way ANOVA, followe by a Tukey's test (*ns,* not significant; *, $p < 0.05$; **, $p < 0.01$; ***, $p < 0.001$; ****, $p < 0.0001$). For eac condition the mean difference, 95% confidence interval (CI) of the difference, and the adjusted *p* valu are shown.

| conditions | two-way ANOVA, followed by Tukey's test | | | |
|---|---|---|---|---|
| **co-localization of Rab5 with GPCR (β-arrestin2)** | | | | |
| basal conditions in Control | **mean diff.** | **95% CI of diff.** | **adjusted *p* value** | |
| b2AR vs. V2b2 | -0.00006492 | -0.4365 to 0.4364 | >0.9999 | ns |
| b2AR vs. b2V2 | 0.103 | -0.3446 to 0.5505 | 0.9336 | ns |
| b2AR vs. V2R | 0.1516 | -0.2784 to 0.5816 | 0.7989 | ns |
| V2b2 vs. b2V2 | 0.103 | -0.3508 to 0.5568 | 0.936 | ns |
| V2b2 vs. V2R | 0.1516 | -0.2848 to 0.5881 | 0.8058 | ns |
| b2V2 vs. V2R | 0.04861 | -0.3989 to 0.4962 | 0.9923 | ns |
| stimulated conditions in Control | | | | |
| b2AR vs. V2b2 | -0.3232 | -0.7539 to 0.1075 | 0.2139 | ns |
| b2AR vs. b2V2 | -0.2593 | -0.6866 to 0.1681 | 0.3983 | ns |
| b2AR vs. V2R | 0.2389 | -0.1822 to 0.66 | 0.459 | ns |
| V2b2 vs. b2V2 | 0.06389 | -0.3757 to 0.5035 | 0.9819 | ns |
| V2b2 vs. V2R | 0.5621 | 0.1286 to 0.9956 | 0.0051 | ** |
| b2V2 vs. V2R | 0.4982 | 0.06804 to 0.9284 | 0.0158 | * |
| basal conditions in ΔQ-GRK | | | | |
| b2AR vs. V2b2 | -0.0000000007 | -0.4326 to 0.4326 | >0.9999 | ns |
| b2AR vs. b2V2 | 0.0000000002 | -0.4131 to 0.4131 | >0.9999 | ns |
| b2AR vs. V2R | -0.0000000004 | -0.4451 to 0.4451 | >0.9999 | ns |
| V2b2 vs. b2V2 | 0.0000000009 | -0.4326 to 0.4326 | >0.9999 | ns |
| V2b2 vs. V2R | 0.0000000003 | -0.4633 to 0.4633 | >0.9999 | ns |
| b2V2 vs. V2R | -0.0000000007 | -0.4451 to 0.4451 | >0.9999 | ns |
| stimulated conditions in ΔQ-GRK | | | | |
| b2AR vs. V2b2 | -0.4035 | -0.8335 to 0.02658 | 0.0748 | ns |
| b2AR vs. b2V2 | -0.2525 | -0.6602 to 0.1551 | 0.3795 | ns |
| b2AR vs. V2R | -0.09571 | -0.5297 to 0.3383 | 0.9409 | ns |
| V2b2 vs. b2V2 | 0.1509 | -0.2791 to 0.581 | 0.8008 | ns |
| V2b2 vs. V2R | 0.3078 | -0.1473 to 0.7628 | 0.3008 | ns |
| b2V2 vs. V2R | 0.1568 | -0.2772 to 0.5908 | 0.7865 | ns |
| basal conditions in ΔQ-GRK + GRK2 | | | | |
| b2AR vs. V2b2 | 0.2988 | -0.2519 to 0.8495 | 0.4989 | ns |
| b2AR vs. b2V2 | -0.01804 | -0.5399 to 0.5039 | 0.9997 | ns |
| b2AR vs. V2R | -0.3651 | -0.9203 to 0.1901 | 0.3257 | ns |
| V2b2 vs. b2V2 | -0.3169 | -0.8512 to 0.2175 | 0.4195 | ns |
| V2b2 vs. V2R | -0.6639 | -1.231 to -0.097 | 0.0143 | * |
| b2V2 vs. V2R | -0.3471 | -0.8861 to 0.1919 | 0.3447 | ns |
| stimulated conditions in ΔQ-GRK + GRK2 | | | | |
| b2AR vs. V2b2 | -0.2824 | -0.8295 to 0.2647 | 0.5421 | ns |
| b2AR vs. b2V2 | -0.1841 | -0.6965 to 0.3283 | 0.7894 | ns |
| b2AR vs. V2R | -0.06903 | -0.6206 to 0.4826 | 0.9883 | ns |
| V2b2 vs. b2V2 | 0.09824 | -0.4306 to 0.6271 | 0.9634 | ns |
| V2b2 vs. V2R | 0.2133 | -0.3536 to 0.7803 | 0.7651 | ns |
| b2V2 vs. V2R | 0.1151 | -0.4184 to 0.6486 | 0.9444 | ns |
| basal conditions in ΔQ-GRK + GRK6 | | | | |
| b2AR vs. V2b2 | 0.2734 | -0.1931 to 0.7399 | 0.43 | ns |
| b2AR vs. b2V2 | 0.1553 | -0.2746 to 0.5851 | 0.7869 | ns |
| b2AR vs. V2R | -0.3518 | -0.8111 to 0.1074 | 0.1981 | ns |
| V2b2 vs. b2V2 | -0.1182 | -0.5592 to 0.3229 | 0.9 | ns |
| V2b2 vs. V2R | -0.6252 | -1.095 to -0.1555 | 0.0037 | ** |
| b2V2 vs. V2R | -0.5071 | -0.9404 to -0.07369 | 0.0144 | * |
| stimulated conditions in ΔQ-GRK + GRK6 | | | | |
| b2AR vs. V2b2 | -0.3074 | -0.7612 to 0.1464 | 0.2998 | ns |
| b2AR vs. b2V2 | 0.5917 | 0.1716 to 1.012 | 0.0018 | ** |
| b2AR vs. V2R | 0.2704 | -0.1798 to 0.7206 | 0.4079 | ns |
| V2b2 vs. b2V2 | 0.8991 | 0.462 to 1.336 | <0.0001 | **** |
| V2b2 vs. V2R | 0.5778 | 0.1117 to 1.044 | 0.0082 | ** |
| b2V2 vs. V2R | -0.3213 | -0.7547 to 0.1121 | 0.2236 | ns |

**Supplementary Table 5:** Detailed statistical results of the analysis of the microscopy co-localization quantification, shown in **Fig. 5b** (for co-localization of Rab5 with β-arrestin2 in Control cells) and **Suppl. Fig. 8**. Here, basal or stimulated values of β-arrestin2 co-localization with Rab5 were compared between tested receptors, using a two-way ANOVA, followed by a Tukey's test (*ns,* not significant; *, $p < 0.05$; **, $p < 0.01$; ***, $p < 0.001$; ****, $p < 0.0001$). For each condition the mean difference, 95% confidence interval (CI) of the difference, and the adjusted *p* value are shown.

| **conditions**<br>**co-localization of Rab5 with β-arrestin2** | **two-way ANOVA, followed by Tukey's test** | | | |
|---|---|---|---|---|
| basal conditions in Control | **mean diff.** | **95% CI of diff.** | **adjusted *p* value** | |
| b2AR vs. V2b2 | -0.04462 | -1.479 to 1.39 | 0.9998 | ns |
| b2AR vs. b2V2 | -1.811 | -3.27 to -0.3522 | 0.0081 | ** |
| b2AR vs. V2R | -0.707 | -2.1 to 0.6859 | 0.556 | ns |
| V2b2 vs. b2V2 | -1.766 | -3.265 to -0.2678 | 0.0135 | * |
| V2b2 vs. V2R | -0.6624 | -2.097 to 0.7721 | 0.6314 | ns |
| b2V2 vs. V2R | 1.104 | -0.3549 to 2.563 | 0.2074 | ns |
| stimulated conditions in Control | | | | |
| b2AR vs. V2b2 | 0.2598 | -1.175 to 1.694 | 0.966 | ns |
| b2AR vs. b2V2 | -9.887 | -11.31 to -8.463 | <0.0001 | **** |
| b2AR vs. V2R | -5.898 | -7.29 to -4.505 | <0.0001 | **** |
| V2b2 vs. b2V2 | -10.15 | -11.61 to -8.682 | <0.0001 | **** |
| V2b2 vs. V2R | -6.157 | -7.592 to -4.723 | <0.0001 | **** |
| b2V2 vs. V2R | 3.989 | 2.566 to 5.412 | <0.0001 | **** |
| basal conditions in ΔQ-GRK | | | | |
| b2AR vs. V2b2 | -0.00000000027 | -0.8105 to 0.8105 | >0.9999 | ns |
| b2AR vs. b2V2 | -0.00000000027 | -0.7793 to 0.7793 | >0.9999 | ns |
| b2AR vs. V2R | -0.00000000027 | -0.8256 to 0.8256 | >0.9999 | ns |
| V2b2 vs. b2V2 | 0 | -0.8156 to 0.8156 | >0.9999 | ns |
| V2b2 vs. V2R | 0 | -0.86 to 0.86 | >0.9999 | ns |
| b2V2 vs. V2R | 0 | -0.8306 to 0.8306 | >0.9999 | ns |
| stimulated conditions in ΔQ-GRK | | | | |
| b2AR vs. V2b2 | -0.02676 | -0.8351 to 0.7816 | 0.9998 | ns |
| b2AR vs. b2V2 | -0.169 | -0.9278 to 0.5897 | 0.9392 | ns |
| b2AR vs. V2R | -1.203 | -2.011 to -0.3944 | 0.0009 | *** |
| V2b2 vs. b2V2 | -0.1423 | -0.9553 to 0.6707 | 0.9691 | ns |
| V2b2 vs. V2R | -1.176 | -2.036 to -0.3165 | 0.0027 | ** |
| b2V2 vs. V2R | -1.034 | -1.847 to -0.2207 | 0.0063 | ** |
| basal conditions in ΔQ-GRK + GRK2 | | | | |
| b2AR vs. V2b2 | -0.006178 | -1.589 to 1.577 | >0.9999 | ns |
| b2AR vs. b2V2 | -1.857 | -3.35 to -0.3631 | 0.008 | ** |
| b2AR vs. V2R | -2.579 | -4.19 to -0.9668 | 0.0003 | *** |
| V2b2 vs. b2V2 | -1.851 | -3.425 to -0.2764 | 0.0138 | * |
| V2b2 vs. V2R | -2.572 | -4.259 to -0.8857 | 0.0006 | *** |
| b2V2 vs. V2R | -0.7218 | -2.325 to 0.8812 | 0.6501 | ns |
| stimulated conditions in ΔQ-GRK + GRK2 | | | | |
| b2AR vs. V2b2 | 0.524 | -1.055 to 2.103 | 0.8267 | ns |
| b2AR vs. b2V2 | -7.5 | -8.986 to -6.014 | <0.0001 | **** |
| b2AR vs. V2R | -4.934 | -6.526 to -3.341 | <0.0001 | **** |
| V2b2 vs. b2V2 | -8.024 | -9.567 to -6.48 | <0.0001 | **** |
| V2b2 vs. V2R | -5.458 | -7.104 to -3.811 | <0.0001 | **** |
| b2V2 vs. V2R | 2.566 | 1.008 to 4.123 | 0.0002 | *** |
| basal conditions in ΔQ-GRK + GRK6 | | | | |
| b2AR vs. V2b2 | -0.06824 | -1.559 to 1.423 | 0.9994 | ns |
| b2AR vs. b2V2 | -1.386 | -2.774 to 0.002559 | 0.0506 | ns |
| b2AR vs. V2R | -3.103 | -4.557 to -1.648 | <0.0001 | **** |
| V2b2 vs. b2V2 | -1.318 | -2.775 to 0.1397 | 0.0923 | ns |
| V2b2 vs. V2R | -3.034 | -4.555 to -1.514 | <0.0001 | **** |
| b2V2 vs. V2R | -1.717 | -3.137 to -0.2964 | 0.0106 | * |
| stimulated conditions in ΔQ-GRK + GRK6 | | | | |
| b2AR vs. V2b2 | 0.467 | -0.9901 to 1.924 | 0.841 | ns |
| b2AR vs. b2V2 | -4.176 | -5.54 to -2.813 | <0.0001 | **** |
| b2AR vs. V2R | -4.827 | -6.273 to -3.382 | <0.0001 | **** |
| V2b2 vs. b2V2 | -4.643 | -6.061 to -3.226 | <0.0001 | **** |
| V2b2 vs. V2R | -5.294 | -6.791 to -3.798 | <0.0001 | **** |
| b2V2 vs. V2R | -0.6509 | -2.056 to 0.7544 | 0.6293 | ns |

**Supplementary Table 6:** Detailed statistical results of the analysis of the microscopy co-localization quantification, shown in **Suppl. Fig. 8**. Here, basal or stimulated values of co-localization for each tested receptor with Rab5, in presence of β-arrestin1, as well as for β-arrestin1 with Rab5 were compared using a two-way ANOVA followed by a Tukey's test (*ns,* not significant; *, $p < 0.05$; **, $p < 0.01$; ***, $p < 0.001$; ****, $p < 0.0001$). For each condition the mean difference, 95% confidence interval (CI) of the difference, and the adjusted *p* value are shown.

| conditions | two-way ANOVA, followed by Tukey's test | | | |
|---|---|---|---|---|
| **co-localization of Rab5 with GPCR (β-arrestin1)** | | | | |
| basal conditions in Control | **mean diff.** | **95% CI of diff.** | **adjusted *p* value** | |
| V2b2 vs. b2V2 | -0.3857 | -0.7531 to -0.01835 | 0.0371 | * |
| V2b2 vs. V2R | -0.5073 | -0.8918 to -0.1228 | 0.0059 | ** |
| b2V2 vs. V2R | -0.1215 | -0.5145 to 0.2714 | 0.7461 | ns |
| stimulated conditions in Control | | | | |
| V2b2 vs. b2V2 | 1.269 | 0.9041 to 1.634 | <0.0001 | **** |
| V2b2 vs. V2R | 0.4831 | 0.1049 to 0.8613 | 0.0081 | ** |
| b2V2 vs. V2R | -0.7859 | -1.17 to -0.4014 | <0.0001 | **** |
| basal conditions in ΔQ-GRK | | | | |
| V2b2 vs. b2V2 | 0.0000000003 | -0.3263 to 0.3263 | >0.9999 | ns |
| V2b2 vs. V2R | 0.0000000006 | -0.3165 to 0.3165 | >0.9999 | ns |
| b2V2 vs. V2R | 0.0000000003 | -0.3218 to 0.3218 | >0.9999 | ns |
| stimulated conditions in ΔQ-GRK | | | | |
| V2b2 vs. b2V2 | 0.1959 | -0.1304 to 0.5222 | 0.3336 | ns |
| V2b2 vs. V2R | 0.02272 | -0.2938 to 0.3393 | 0.9843 | ns |
| b2V2 vs. V2R | -0.1732 | -0.4949 to 0.1486 | 0.4131 | ns |
| basal conditions in ΔQ-GRK + GRK2 | | | | |
| V2b2 vs. b2V2 | 0.01026 | -0.2981 to 0.3186 | 0.9966 | ns |
| V2b2 vs. V2R | -0.6851 | -0.9934 to -0.3767 | <0.0001 | **** |
| b2V2 vs. V2R | -0.6953 | -1.014 to -0.3769 | <0.0001 | **** |
| stimulated conditions in ΔQ-GRK + GRK2 | | | | |
| V2b2 vs. b2V2 | 1.161 | 0.863 to 1.459 | <0.0001 | **** |
| V2b2 vs. V2R | 0.1548 | -0.1501 to 0.4597 | 0.4555 | ns |
| b2V2 vs. V2R | -1.006 | -1.319 to -0.6943 | <0.0001 | **** |
| basal conditions in ΔQ-GRK + GRK6 | | | | |
| V2b2 vs. b2V2 | 0.2773 | -0.1073 to 0.6619 | 0.2067 | ns |
| V2b2 vs. V2R | -0.7664 | -1.136 to -0.3971 | <0.0001 | **** |
| b2V2 vs. V2R | -1.044 | -1.426 to -0.6617 | <0.0001 | **** |
| stimulated conditions in ΔQ-GRK + GRK6 | | | | |
| V2b2 vs. b2V2 | 1.37 | 0.9935 to 1.745 | <0.0001 | **** |
| V2b2 vs. V2R | 0.3362 | -0.02763 to 0.7001 | 0.0767 | ns |
| b2V2 vs. V2R | -1.033 | -1.412 to -0.6548 | <0.0001 | **** |
| **co-localization of Rab5 with β-arrestin1** | | | | |
| basal conditions in Control | **mean diff.** | **95% CI of diff.** | **adjusted *p* value** | |
| V2b2 vs. b2V2 | 0.243 | -1.419 to 1.905 | 0.9365 | ns |
| V2b2 vs. V2R | -2.015 | -3.715 to -0.3153 | 0.0154 | * |
| b2V2 vs. V2R | -2.258 | -4.017 to -0.4999 | 0.0077 | ** |
| stimulated conditions in Control | | | | |
| V2b2 vs. b2V2 | -6.228 | -7.923 to -4.532 | <0.0001 | **** |
| V2b2 vs. V2R | -10.38 | -12.09 to -8.675 | <0.0001 | **** |
| b2V2 vs. V2R | -4.156 | -5.938 to -2.374 | <0.0001 | **** |
| basal conditions in ΔQ-GRK | | | | |
| V2b2 vs. b2V2 | 0.2149 | -0.4663 to 0.8961 | 0.7368 | ns |
| V2b2 vs. V2R | 0 | -0.6552 to 0.6552 | >0.9999 | ns |
| b2V2 vs. V2R | -0.2149 | -0.8868 to 0.457 | 0.7306 | ns |
| stimulated conditions in ΔQ-GRK | | | | |
| V2b2 vs. b2V2 | 0.1311 | -0.5562 to 0.8185 | 0.8942 | ns |
| V2b2 vs. V2R | -0.5662 | -1.221 to 0.08908 | 0.1053 | ns |
| b2V2 vs. V2R | -0.6973 | -1.375 to -0.01918 | 0.0423 | * |
| basal conditions in ΔQ-GRK + GRK2 | | | | |
| V2b2 vs. b2V2 | -0.3451 | -1.957 to 1.267 | 0.8688 | ns |
| V2b2 vs. V2R | -1.386 | -2.997 to 0.2261 | 0.1077 | ns |
| b2V2 vs. V2R | -1.041 | -2.715 to 0.6335 | 0.3089 | ns |
| stimulated conditions in ΔQ-GRK + GRK2 | | | | |
| V2b2 vs. b2V2 | -7.985 | -9.629 to -6.342 | <0.0001 | **** |
| V2b2 vs. V2R | -9.887 | -11.5 to -8.271 | <0.0001 | **** |
| b2V2 vs. V2R | -1.901 | -3.626 to -0.1765 | 0.0267 | * |
| basal conditions in ΔQ-GRK + GRK6 | | | | |
| V2b2 vs. b2V2 | 0.1246 | -1.298 to 1.547 | 0.9767 | ns |
| V2b2 vs. V2R | -1.823 | -3.235 to -0.4118 | 0.0073 | ** |
| b2V2 vs. V2R | -1.948 | -3.38 to -0.5155 | 0.0044 | ** |
| stimulated conditions in ΔQ-GRK + GRK6 | | | | |
| V2b2 vs. b2V2 | 0.4999 | -0.9585 to 1.958 | 0.6975 | ns |
| V2b2 vs. V2R | -7.39 | -8.801 to -5.978 | <0.0001 | **** |
| b2V2 vs. V2R | -7.89 | -9.358 to -6.421 | <0.0001 | **** |
